# *In-Vivo* Toxicity Studies and *In-Vitro* Inactivation of SARS-CoV-2 by Povidone-iodine *In-situ* Gel Forming Formulations

**DOI:** 10.1101/2020.05.18.103184

**Authors:** Bo Liang, Xudong Yuan, Gang Wei, Wei Wang, Ming Zhang, Haizhou Peng, Amin Javer, Michelle Mendenhall, Justin Julander, Sammi Huang, Hany Michail, Yong Lu, Qi Zhu, John Baldwin

**Affiliations:** IView Therapeutics, Doylestown, PA 18902, USA (B. Liang, X. Yuan, W. Wang, M. Zhang, S. Huang, H. Michail, Q. Zhu, J. Baldwin); School of Pharmacy, Fudan University, Shanghai, China (G. Wei); IVIEW Therapeutics (Zhuhai) Co. Ltd. Hengqin, Zhuhai, China (B. Liang, X. Yuan, W. Wang, H. Peng, Y. Lu, Q. Zhu); St. Paul’s Sinus Centre, University of British Columbia, Vancouver, Canada (A. Javer); Institute for Antiviral Research, Utah State University, Logan, UT 84322, USA (M. Mendenhall, J. Julander)

**Keywords:** COVID-19, SARS-CoV-2, coronavirus, nasal spray, eye drop, viral inactivation, infection prevention, Povidone-iodine, PVP-I, in-situ gel, sustained release

## Abstract

To curb the spread of SARS-CoV-2, the etiologic agent of the COVID-19 pandemic, we characterize the virucidal activity of long-acting Povidone Iodine (PVP-I) compositions developed using an *in-situ* gel forming technology. The PVP-I gel forming nasal spray (IVIEW-1503) and PVP-I gel forming ophthalmic eye drop (IVIEW-1201) rapidly inactivated SARS-CoV-2, inhibiting the viral infection of VERO76 cells. No toxicity was observed for the PVP-I formulations. Significant inactivation was noted with preincubation of the virus with these PVP-I formulations at the lowest concentrations tested. It has been demonstrated that both PVP-I formulations can inactivate SARS-CoV-2 virus efficiently in both a dose-dependent and a time-dependent manner. These results suggest IVIEW-1503 and IVIEW-1201 could be potential agents to reduce or prevent the transmission of the virus through the nasal cavity and the eye, respectively. Further studies are needed to clinically evaluate these formulations in early-stage COVID-19 patients.

## Introduction

The ongoing coronavirus pandemic known as coronavirus disease 2019 (COVID-19), is caused by severe acute respiratory syndrome coronavirus 2 (SARS-CoV-2) (1). The outbreak was first identified in Wuhan, Hubei, China, in December 2019, and was recognized as a pandemic by the World Health Organization (WHO) on March 11 2020 (2). According to Johns Hopkins dashboard, as of May 18, 2020, there are over 4.8 million confirmed cases of COVID-19 that have been reported in 188 countries and territories and claimed more than 318,000 lives. More than 1.5 million cases were confirmed in the U.S.

Coronaviruses are a group of viruses that cause a significant percentage of all common colds in human adults and children. Four human coronavirus including 229E, OC43, NL63, and HKU1 are prevalent and typically cause common cold symptoms in immunocompetent individuals (3). Severe acute respiratory syndrome coronavirus 2 (SARS-CoV-2) is the third coronavirus to cross species to infect human populations (probably transmitted from bats or another host), causing pandemics over the past two decades (4,5,6). The previous two were the severe acute respiratory syndrome coronavirus (SARS-CoV) outbreak in 2002 and the Middle East respiratory syndrome coronavirus (MERS-CoV) outbreak in 2012 (7,8). SARS-Cov-2 has a unique pathogenesis because it causes both upper and lower respiratory tract infections (9). SARS-CoV-2 is classified as a novel betacoronavirus belonging to the sarbecovirus subgenus of Coronaviridae family. The genome sequence of SARS-CoV-2 is about 89% identical to bat SARS-like-CoVZXC21 and 82% identical to human SARS-CoV. It has been reported that SARS-CoV-2 uses the same cell entry receptor, ACE2, to infect humans as SARS-CoV, so clinical similarity between the two viruses is expected, particularly in severe cases (10).

The virus is typically spread by close physical contact and via respiratory droplets when people cough or sneeze (11,12,13). People may also catch COVID-19 by touching their eyes, nose, or mouth immediately after touching a contaminated surface (12,13). It is most contagious when people are symptomatic, however, it may also spread when a person is asymptomatic (13). Nasal cells are identified as the key entry point for SARS-CoV-2 (14). Goblet and ciliated cells in the nose have high levels of the entry proteins ACE-2 and TMPRSS2 that SARS-CoV-2 uses to get into human cells. The virus exploits existing secretory pathways in nasal goblet cells sustained at a pre-symptomatic stage. Nasal carriage is likely to be a key feature of transmission, therefore drugs administered intranasally could be highly effective in limiting spread. The two key entry proteins ACE2 and TMPRSS2 were also found in cells in the cornea of the eye, ocular as another route of infection (14, 15). With SARS-CoV-2 detected in human tears, ocular secretions might be a potential for human to human spread (16). There is controversy on whether the eye could be a transmission route for COVID-19, however, even if COVID 19 positive patients have no virus shedding through tears, the eye remains a potential portal of entry for ocular and systemic disease from the virus (16). Therefore, developing nasal and ocular treatment to reduce and prevent transmission of the debilitating virus is a viable while potentially highly effective strategy.

There is no known vaccine or specific antiviral treatment for COVID-19, but development efforts are underway, including testing of existing medications (12). Current primary treatment is limited to supportive treatment of symptoms. Recommended preventive measures include hand washing, covering the mouth when coughing, maintaining distance from other people, and monitoring and self-isolation for people suspected for being infected (12). The COVID-19 case fatality rate has not yet peaked and it is anticipated that many more deaths will occur despite the efforts from many countries. There is, therefore, an urgent need to find effective ways to prevent and treat this virus.

Povidone-iodine (PVP-I) is a complex of polyvinylpyrrolidone and iodine. It is also called iodophor and contains 9-12% iodine. It is a powerful disinfectant with broad-spectrum application and is effective against viruses, bacteria, fungi, and mold spores. PVP-I products have been used as disinfectant for the inactivation of various bacteria and viruses for years because of their strong bactericidal and antiviral activities. Povidone-iodine (PVP-I) is routinely used in ophthalmology and general surgery. PVP-I has been used in acute and chronic treatment of a variety of human indications. There have been numerous clinical studies demonstrating the safety of PVP-I in a variety of topical applications in ophthalmology, otology, rhinology and dermatology (18,19,20,21,22,23,24). PVP-I in nasal usage was also well documented, including both single use (24,25) and multiple applications. 3M Company introduced PVP-I based skin and nasal antiseptic (Povidone Iodine Solution 5% w/w [0.5% available iodine] USP). It has rapid, broad-spectrum antimicrobial activity, which reduces nasal bacteria, including *S. aureus*, by 99.5% in just one hour and maintains this reduction for at least 12 hours (26,27). In the clinical setting, rhinologists and oral surgeons commonly use higher concentrations of PVP-I for various antimicrobial therapies as it offers an affordable, potent, well-tolerated and widely available antiseptic option with little to no cross-resistance. Gluck and colleagues reported a phase I study (n=35) assessing tolerability and local effect of a PVP-I spray (PVP-I 2.2 or 4.4%) in single and repeated (3 times a day for 3 days). No safety-relevant finding or serious adverse events were reported, no evidence for cyto-nor genotoxicity obtained (28).

It was reported that PVP-I formulations are effective against both enveloped and non-enveloped viruses (29, 30, 31). The virucidal activity is mainly due to the free iodine released from PVP-I (32). In one study, PVP-I was effective in inhibiting SARS-CoV infectivity (33) of SARS-CoV with PVP-I products reduced the virus infectivity after 2 minutes of exposure from 1.17 x 10^6^ TCID (50)/ml to below the detectable level. In another study, PVP-I gargle/mouthwash diluted 1:30 (equivalent to a concentration of 0.23% PVP-I) rapidly inactivated SARS-CoV after 15 seconds of exposure (34). An *in-vitro* study of three formulations of PVP-I (PVP-I: 4% PVP-I skin cleanser, 7.5% PVP-I surgical scrub, and 1% PVP-I gargle/mouthwash) against a reference virus (Modified vaccinia virus Ankara, MVA) and MERS-CoV (35) showed a reduction in virus titer of ≥4 Log10 versus MVA and MERS-CoV, under both clean and dirty conditions after 15 seconds of exposure with each undiluted PVP-I product. In addition to antiviral activity of SARS-CoV and MERS-CoV, PVP-I also showed virucidal efficacy against influenza virus A (H1N1), rotavirus, and murine norovirus (MNV) (34, 36, 37). PVP-I has also shown efficacy in preventing the infection and limiting the spread of Ebola virus disease (EBV) (38). Additionally, it is reported that PVP-I is virucidal against non-enveloped viruses e.g. polyomavirus, adenovirus, and poliovirus type 1 (31). Due to the similarities among enveloped viruses, PVP-I is expected to show virucidal efficacy against SARS-CoV-2. However, there is no published data on the use of iodine for the treatment or prevention of COVID-19. Currently no PVP-I product has been approved as a potential protection and prevention for the spread of this new debilitating virus. UK investigators recommended the immediate and UK-wide use of PVP-I nasal spray and mouthwash in healthcare workers and their patients to minimize the risk of spread of COVID-19 as an adjunct to currently recommended PPE (39).

It’s well known that nasal solutions are cleared off rapidly from the nasal cavity, while the conventional liquid ocular formulation is eliminated from the precorneal area immediately upon instillation because of lacrimation and effective nasolacrimal drainage (40). The short contact time of water-soluble povidone-iodine on nasal mucosa or in the eye is undesirable and would necessitate frequent and multiple administration to maintain the virucidal efficacy, thereby limiting practicality and increasing medical burden for the patient. In addition, frequent dosing can lead to irritation and potential toxicity. Therefore, developing a safe, non-toxic and long-acting PVP-I nasal spray or ophthalmic eye drop is an urgent medical need to protect people from SARS-CoV-2 infection, as well as to block the transmission through the nasal cavity or through the eye.

In the current study, we investigated the *in-vitro* virucidal efficacy of PVP-I *in-situ* gel forming formulations against SARS-CoV-2 virus along with their safety assessment in animal toxicology studies. The results suggest that both IVIEW-1201 and IVIEW-1503, reduced SARS-CoV-2 viral titers to near or below the level of detection after 2 minutes of exposure. Therefore, these preparations could potentially be used as a disinfectant for the virus in the nasal cavity or in the eye, respectively. By employing either the sustained release nasal spray or ophthalmic eye drop delivery technologies, we expect to reduce, treat, and eliminate SARS-CoV-2 in the nasal, sinus cavity, and in the eye. Additionally, the long-acting nasal spray can potentially be utilized as a prophylaxis for protecting against SARS-CoV-2 infection.

## Materials and Methods

### Chemicals

Two sustained release povidone-iodine *in-situ* gel forming formulations, IVIEW-1201 and IVIEW-1503, consisting of 1.0 % and 0.6% of Povidone-iodine (w/w), respectively, were prepared for the study. Simulated tear fluid consisted of NaHCO_3_ (CAS 144-55-8), NaCl (CAS 7647-14-5), CaCl_2_ (CAS 7440-70-2), KCl (CAS 7447-40-7) (Spectrum Chemicals, New Brunswick, NJ) and water. Simulated nasal fluid consisted of NaCl, CaCl_2_, KCl and water. In addition, ethanol (CAS 64-17-5) (Spectrum Chemicals, New Brunswick, NJ) was used as supplied. Povidone-iodine (1-vinyl-2-pyrrolidinone polymers, iodine complex], USP, CAS 25655-41-8) is supplied by Ashland.

### Virus strains and cell culture

SARS-CoV-2 (strain USA_WA1/2020, prepared by Natalie Thornburg, CDC and provided by WRCEVA, University of Texas Medical Branch) virus stocks were prepared by growing virus in VERO 76 cells (ATCC^®^, CRL-1587, ATCC, Manassas, Virginia). Test media used was MEM supplemented with 2% fetal bovine serum (FBS, GE Healthcare Hyclone, Marlborough, MA) and 50 μg/mL gentamicin (Sigma-Aldrich, St. Louis, Missouri).

### Formulation Development

Both formulations (IVIEW-1201 and IVIEW-1503) are brownish aqueous gel forming solutions of containing 1.0% and 0.6% (w/w) polyvinylpyrrolidinone-iodine complex (Povidone-Iodine), [1-vinyl-2-pyrrolidinone polymers, iodine complex], respectively. IVIEW-1201 is packaged in polypropylene (PP) plastic eye drop bottle; and IVIEW-1503 is packaged in amber glass bottle equipped with Aptar nasal spray pump for intranasal application.

We have developed a proprietary PVP-I based sustained release platform technology (41,42) to produce *in-situ* gel forming formulations where the effective concentration of PVP-I is maintained by the equilibrium between solution PVP-I and the gel bound components resulting in a long-lasting, less toxic pharmacological effect in the eye and nasal cavity. The *in-situ* gel forming PVP-I composition is formulated with ion-sensitive *in-situ* gel forming materials such as Deacetylated Gellan Gum (Gelrite^®^) to increase the residence time of the dosage form in the eye and on the nasal mucosa. Preparations of PVP-I with gellan gum are dropped into eyes or sprayed into nasal cavity; gel formation takes place, induced by the electrolytes (Na^+^, K^+^, Ca^2+^, etc.) of the tear fluid or nasal fluid (43).

## Toxicological Studies of Dilute PVP-I Formulations

### Ocular Toxicity Study

We have conducted a 7-Day Repeat Topical Ocular Dose Toxicity Study in rabbits to determine the potential ocular toxicity of PVP-I gel forming formulations at two concentrations, and a PVP-I /dexamethasone (dex) formulation (PVP-I 0.6%/dex 0.1%, this formulation was advanced by Shire Llc. for phase III clinical trials against adenoviral conjunctivitis in 2017 as SHP640) comparing each to the vehicle when administered by repeat topical ocular doses over a 7-day period to New Zealand white rabbits. Only rabbits with clinical normal eyes were used in this study. 12 female rabbits were equally divided into four groups with three (3) rabbits per group. Group one is vehicle, Group 2 is test group using 0.6% (6 mg/mL) PVP-I gel forming solution. Group 3 is a test group using 1.0% (10 mg/mL) PVP-I gel forming solution, and Group 4 is a test group using PVP-I 0.6% (6 mg/mL)/ dex 0.1% (1 mg/mL) suspension. Each eye was dosed 35 μL/eye/dose for twice a day.

Ophthalmic examinations occurred pre-dose, Day 1 after the first dose and after the final daily dose, Day 4 (±1 day) after the final daily dose, and on Day 7 after the final daily dose, but prior to necropsy. The evaluations included slit lamp examination, including general scoring according the McDonald-Shadduck scoring scheme, assessments of fluorescein and rose Bengal staining according to the National Eye Institute (NEI) corneal staining grid (5 sectors assessed for type of staining, surface area, and density of staining), and an assessment of eyelid swelling. In addition, corneal sensation was assessed via a Cochet-Bonnet esthesiometer. At the conclusion of the study, the animals were humanely euthanized, and the eyes and eyelids assessed microscopically. The fluorescein and rose Bengal scores were assessed by calculating a global score for each eye at each time point as well as score incidence.

### Intranasal Administration Toxicity Study

We have conducted a 28-day toxicity study using Sprague Dawley CD^®^ IGS rats to determine the potential subchronic toxicity of a PVP-I formulation (0.8% PVP-I/0.064% Budesonide gel forming nasal spray formulation). Sixty healthy rats (60) were selected for the test and equally distributed into four test groups and two recovery groups (control and high dose). Intranasal administration of the formulation at dose levels of 25, 50 and 75 μl and saline control at dose levels of 75 μl were evaluated. The saline control or the test substance was administered into the right nostril via a 200 μl pipette twice daily (approximately 12 hours apart). The animals were observed at least once daily for viability, signs of gross toxicity, and behavioral changes, and weekly for a battery of detailed observations. All main study animals were subjected to a necropsy of the upper respiratory tract and related sinuses at study termination (Day 29). Thyroids and lungs were collected and weighed.

This formulation has higher PVP-I concentration compared to IVIEW-1503 formulation (0.6%). Since the intranasal usage of high concentration of PVP-I had no safety issues in the subchronic toxicity study, IVIEW-1503 as a nasal spray formulation should also have minimal to no safety concern.

### Virucidal Assay

The IVIEW-1201 (1.0% PVP-I) and IVIEW-1503 (0.6% PVP-I) formulations were tested for virucidal activity at the following concentrations: full strength (90% sample and 10% virus solution), 1/1.8, 1/3.2, and 1/10 diluted in simulated tears or simulated nasal fluid, respectively.

SARS-CoV-2 virus stock was added to triplicate tubes of each prepared concentration at 1/10, so the final concentrations of solution tested were 90%, 50%, 28% and 9% of the original formulation concentration. Thus, the final PVP-I concentrations for IVIEW-1201 are 0.90%, 0.50%, 0.28%, and 0.09%. The final PVP-I concentrations for IVIEW-1503 are 0.54%, 0.30%, 0.17%, and 0.05%. Tear or nasal fluid only was added to one tube of each prepared concentration in the presence of virus to serve as the toxicity control. Ethanol (45%) was tested in parallel as the positive control and water only to serve as the virus control.

Solution and virus were incubated at 37°C for three contact times of 30 seconds, 2 minutes, and 10 minutes. Following the contact period, the solutions were neutralized by a 1/10 dilution in test media containing 10% FBS and 0.5% sodium thiosulfate.

### Virus Quantification

Neutralized samples were serially diluted using eight half-log dilutions in the test medium. Each dilution was added to 4 wells of a 96-well plate with 80-100% confluent VERO 76 cells. The toxicity controls were added to an additional 4 wells and 2 of these wells were infected with virus to serve as neutralization controls, ensuring that the neutralized samples did not continue to inhibit growth and detection of surviving virus. All plates were incubated at 37°C, 5% CO_2_.

On day 6, the post-infection plates were scored for presence or absence of viral cytopathic effect (CPE). The Reed-Muench method was used to determine end-point titers (50% cell culture infectious dose, CCID50) of the samples, and the log reduction value (LRV) of the compound compared to the negative (water) control was calculated.

### Controls

The reduction of virus in formulation-treated test wells compared to virus controls was calculated as the log reduction value (LRV). Toxicity controls were tested with media not containing virus to see if the samples were toxic to cells. Neutralization controls were tested to ensure that virus inactivation did not continue after the specified contact time, and that residual sample in the titer assay plates did not inhibit growth and detection of surviving virus. This was done by adding toxicity samples to titer test plates then spiking each well with a low amount of virus (~60 CCID50) that would produce an observable amount of CPE during the incubation period.

## Results

### Ocular Toxicity Study Results

No meaningful signs of ocular irritation or corneal staining were observed in the vehicle group (Group 1) during the study. Repeated topical dosing of each povidone iodine-containing formulation was associated with brief behavioral signs of irritation after each drop (squinting and pawing of eyes), transient signs of mild ocular irritation, and persistent corneal staining.

In Groups 2 and 3, mild to moderate ocular irritation was observed, characterized primarily by conjunctival involvement, and was more pronounced on Day 1 after the last dose, then lessened with time. A dose response with the irritation was observed whereby Group 3 (1.0% PVP-I) showed more signs of irritation, including transient corneal involvement, than Group 2 (0.6% PVP-I). At the end of Day 1, the mean global staining score was highest for Group 2 (0.6% PVP-I) for both fluorescein and rose Bengal staining as compared to Group 3 (1.0% PVP-I). The corneal staining in Group 2 (0.6% PVP-I) was lower at the Day 4 exam; the fluorescein staining plateaued between Days 4 and 7, whereas the rose Bengal staining continued to decrease in intensity. On Days 4 and 7, the staining in Group 2(0.6% PVP-I) was lower than the staining for Groups 3 (1.0% PVP-I). The fluorescein and rose Bengal staining scores for Groups 3 (1.0% PVP-I) did not change dramatically from Day 1 last dose scores.

In conclusion, the repeated topical administration of povidone iodine-containing gel forming formulations over 7-days was associated with mild and transient signs of ocular irritation. Corneal staining with both fluorescein and rose Bengal was observed in both Group 2 (0.6% PVP-I) and Group 3 (1.0% PVP-I). After the last dose on Day 1, the Group 2 (0.6% PVP-I) mean global staining score was higher than Group 3 (1.0% PVP-I). However, the staining in Group 2 (0.6% PVP-I) decreased by Day 4 such that the scores were lower than the staining in Group 3 (1.0% PVP-I), which changed little from the Day 1 observations. There were no associated histological findings. The results suggest that the 0.6% PVP-I and 1.0% PVP-I gel formulations (Group 2 and 3) are associated with transient irritation and mild toxicity limited to the superficial ocular tissues with a slight dose response suggested after Day 1. The lack of meaningful histological signs of corneal damage or inflammation support that the clinical observations are not associated with permanent changes after 7 days of dose administration.

### Intranasal Administration Toxicity Study Results

There were no mortalities during the course of the study and no test substance-related changes in body weight, body weight gain, food consumption, thyroid weights and lung weights for the duration of the study. There were no macroscopic observations at terminal sacrifice considered attributable to the administration of the formulation intranasally. Following a gross necropsy of the upper respiratory tract and related sinuses, there were no signs of irritation and no abnormalities were detected. Therefore, it has demonstrated the subchronic intranasal use of PVP-I is expected to have no toxicity concerns.

### Virucidal Assay Results

Virus LRV (log reduction value) for IVIEW-1201 and IVIEW-1503 against SARS-CoV-2 virus are shown in **Table 1**. PVP-I formulation toxicity was not observed at any concentration. Ethanol (45%) had some observable toxicity at the 30-second and 2-minute time points. As a result of this toxicity, the presence of virus could not be ruled out in those wells therefore the limit of detection was 1.7 log_10_ CCID_50_ of virus per 0.1 mL.

**Table 1.** Virucidal efficacy against Covid-19 virus after incubation with virus at 37°C.

| Sample | Drug | PVP-I | Contact | Viral Titer<sup>a</sup> | LRV<sup>b</sup> |
| --- | --- | --- | --- | --- | --- |
|  | Concentration (%) | Concentration (%) | Time (min) |  |  |
| IVIEW-1201 | 90 | 0.9 | 0.5 | <0.67 ± 0.0 | 3.5 |
| IVIEW-1201 | 50 | 0.5 | 0.5 | 1.0 ± 0.3 | 3.2 |
| IVIEW-1201 | 28 | 0.28 | 0.5 | 2.0 ± 1.0 | 2.2 |
| IVIEW-1201 | 9 | 0.09 | 0.5 | 3.0 ± 0.4 | 1.2 |
| IVIEW-1503 | 90 | 0.54 | 0.5 | 1.1 ± 0.1 | 3.1 |
| IVIEW-1503 | 50 | 0.3 | 0.5 | 1.1 ± 0.3 | 3.1 |
| IVIEW-1503 | 28 | 0.17 | 0.5 | 1.2 ± 0.4 | 2.9 |
| IVIEW-1503 | 9 | 0.05 | 0.5 | $1.9 \pm 1.0$ | 2.3 |
| Ethanol | | | 0.5 | $<1.7 \pm 0.0$ | 3.5 |
| Virus Control | | | 0.5 | $4.2 \pm 0.4$ | 0 |
| IVIEW-1201 | 90 | 0.9 | 2 | $<0.67 \pm 0.0$ | 2.9 |
| IVIEW-1201 | 50 | 0.5 | 2 | $<0.67 \pm 0.0$ | 2.9 |
| IVIEW-1201 | 28 | 0.28 | 2 | $1.0 \pm 0.3$ | 2.6 |
| IVIEW-1201 | 9 | 0.09 | 2 | $2.8 \pm 0.4$ | 0.8 |
| IVIEW-1503 | 90 | 0.54 | 2 | $<0.67 \pm 0.0$ | 2.9 |
| IVIEW-1503 | 50 | 0.3 | 2 | $<0.67 \pm 0.0$ | 2.9 |
| IVIEW-1503 | 28 | 0.17 | 2 | $0.8 \pm 0.2$ | 2.8 |
| IVIEW-1503 | 9 | 0.05 | 2 | $1.7 \pm 0.6$ | 1.9 |
| Ethanol | | | 2 | $<1.7 \pm 0.0$ | 1.9 |
| Virus Control | | | 2 | $3.6 \pm 0.1$ | 0 |
| IVIEW-1201 | 90 | 0.9 | 10 | $<0.67 \pm 0.0$ | 3.3 |
| IVIEW-1201 | 50 | 0.5 | 10 | $<0.67 \pm 0.0$ | 3.3 |
| IVIEW-1201 | 28 | 0.28 | 10 | $<0.67 \pm 0.0$ | 3.3 |
| IVIEW-1201 | 9 | 0.09 | 10 | $2.9 \pm 0.4$ | 1 |
| IVIEW-1503 | 90 | 0.54 | 10 | $<0.67 \pm 0.0$ | 3.3 |
| IVIEW-1503 | 50 | 0.3 | 10 | $<0.67 \pm 0.0$ | 3.3 |
| IVIEW-1503 | 28 | 0.17 | 10 | $<0.67 \pm 0.0$ | 3.3 |
| IVIEW-1503 | 9 | 0.05 | 10 | $2.4 \pm 0.1$ | 1.6 |
| Ethanol | | | 10 | $<0.67 \pm 0.0$ | 3.3 |
| Virus Control | | | 10 | $4.0 \pm 0.3$ | 0 |

<sup>a</sup>: Log10 CCID50 of virus per mL, mean of 3 replicates ± standard deviation
<sup>b</sup>: LRV (log reduction value) is the reduction of virus compared to the virus control

In antiviral kinetics studies, a dose response was observed after treatment with both IVIEW-1201 (**Figure 1**) and IVIEW-1503 (**Figure 2**). PVP-I formulations produced greater reduction in virus with increasing concentration and time of contact with the virus. Higher concentrations (0.9% PVP-I of IVIEW-1201; 0.54% PVP-I of IVIEW-1503) of the formulations completely inactivated SARS-CoV-2 virus, reducing titers below the level of detection. This was similar for the half concentration of both IVIEW-1201 (0.5%) and IVIEW-1503 (0.3%), which also reduced virus to near or below the level of detection. Lower concentrations (0.28% PVP-I of IVIEW-1201; 0.17% PVP-I of IVIEW-1503) also reduced the virus substantially. The lowest concentration (0.09% PVP-I of IVIEW-1201; 0.05% PVP-I of IVIEW-1503) of the formulations did not reduce virus significantly with increased contact time.

**Figure 1.**
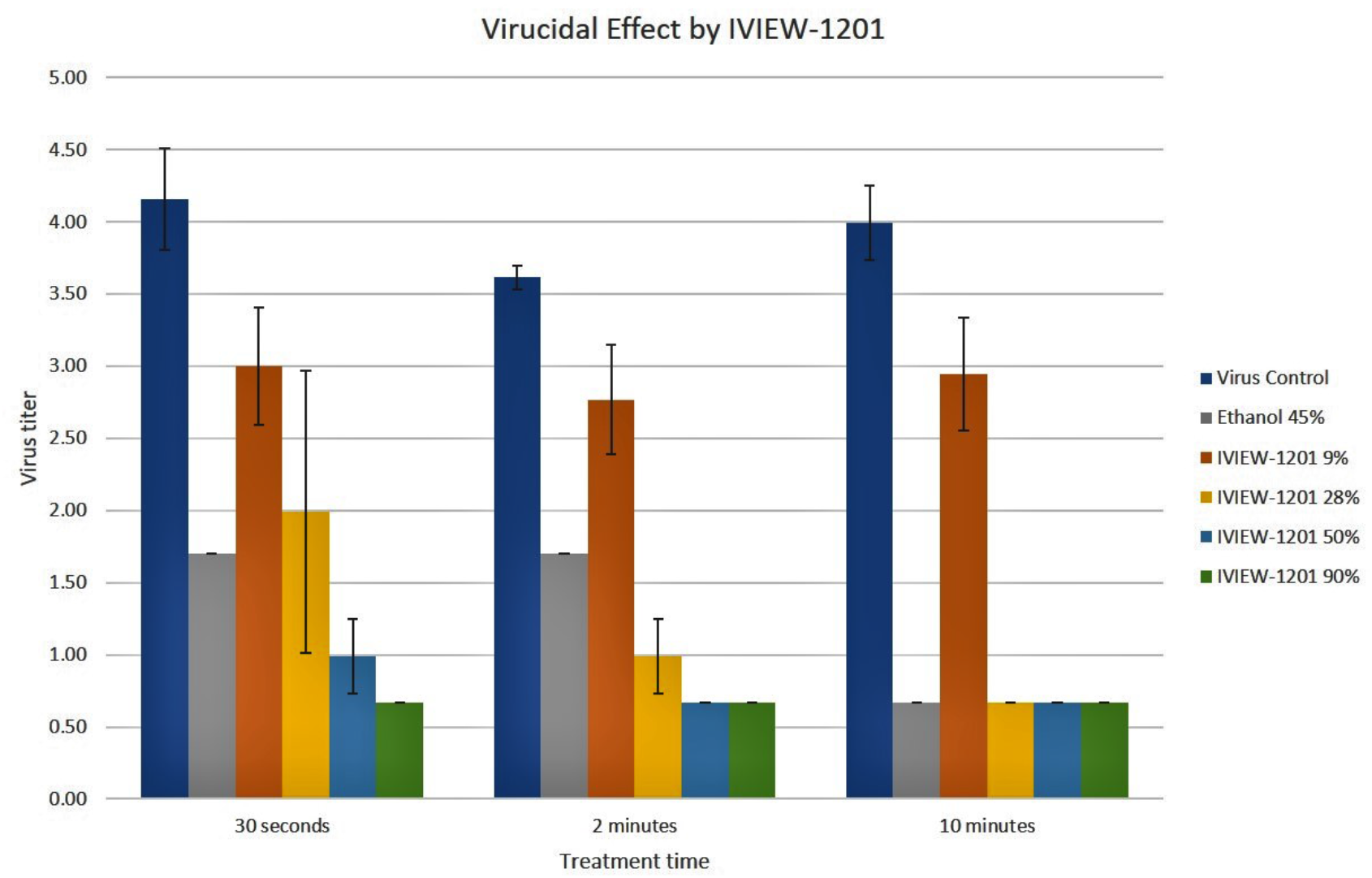
Kinetics of SARS-CoV-2 Virus Inactivation by IVIEW-1201 PVP-I Gel Forming Ophthalmic Formulation

**Figure 2.**
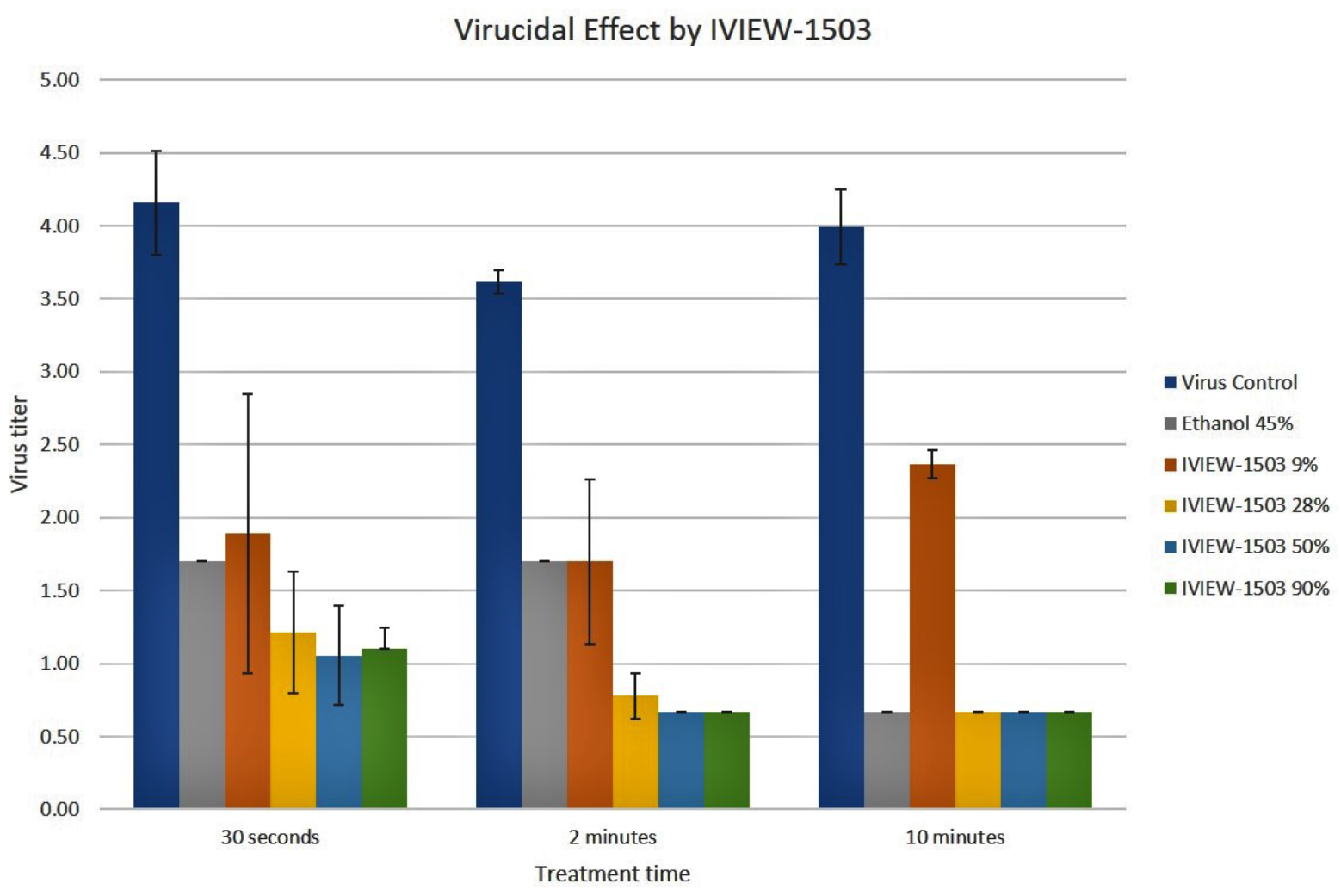
Kinetics of SARS-CoV-2 Virus Inactivation by IVIEW-1503 PVP-I Gel-Forming Nasal Spray Formulation

Neutralization controls demonstrated that residual samples did not inhibit virus growth and detection in the endpoint titer assays in wells that did not have cytotoxicity. Virus controls and positive controls performed as expected.

## Discussion

Although several potential treatments have been identified, there are currently no clinically approved interventions for the treatment or prevention of SARS-CoV-2 virus. A variety of drugs such as Interferon α, Lopinavir/Ritonavir, Ribavirin, Chloroquine Phosphate and Arbidol have undergone limited clinical evaluation in China, U.S. and Italy, but do not show a clear benefit to patients (44, 45, 46, 47, 48). One of the major limitations of the drugs mentioned above is that there is no data available on the adverse reactions of the high dosages of the antiviral drugs when given for long durations. The duration of medication needed is also currently unknown (48).

Studies have shown that the SARS-CoV-2 virus uses angiotensin-converting enzyme 2 (ACE2) in the lungs as a receptor and relies on an enzyme known as transmembrane protease serine 2 (TMPRSS2) to enter and fuse with host cells in order to replicate (49). This has led medical professionals to test existing antivirals that are known to interfere with viral membrane fusion and disrupt RNA polymerase in order to prevent further viral production in hosts. Hydroxychloroquine, an antimalarial and derivative of chloroquine, is one such therapy that is believed to be one of the potential therapies in treating COVID-19. Its mechanism has been used to prevent the transport of virions to the replication site by altering pH and block viral endocytosis to hosts (49). However, in a recent COVID-19 trial consisting of 36 patients, 14 patients treated with hydroxychloroquine have continued to show symptoms of upper or lower respiratory tract infections (50). Although hydroxychloroquine has increased potency and tolerable safety profile compared to chloroquine, potential adverse effects such as electrolyte imbalance and fatal dysrhythmias could take place (50). Another possible therapy, Remdesivir^®^, a nucleotide analogue, inhibits viral RNA polymerase and was developed as a substance to combat ebolaviruses, SARS-CoV, and MERS-CoV (49, 52, 53). With significant results in *in vitro* testing against SARS-CoV-2, the therapy appears to be another potential prophylaxis. In a compassionate use study of 53 COVID-19 patients, 36 patients (68%) have showed clinical improvement in the category of oxygen support while 8 patients (15%) showed worsening (52, 53). Remdesivir^®^ showed significant improvement among patients with minimal to no invasive oxygen support. Remdesivir^®^ is more effective in patients who are not receiving invasive ventilation or extracorporeal membrane oxygenation (ECMO) (52, 53). During the study, there have been no new safety concerns, however, the relatively short duration and small size study does not provide clear evidence of toxicity (52, 53). Nonetheless, FDA has approved the use of hydroxychloroquine and Remdesivir^®^ for emergency authorization to treat COVID-19.

We have successfully developed the *in-situ* gel ophthalmic formulation, IVIEW-1201, currently in global phase II trials for the treatment of adenoviral conjunctivitis; and an *in-situ* gel forming PVP-I/Budesonide combination nasal spray formulation, IVIEW-1502, under pre-IND studies for the treatment of chronic rhinosinusitis. The development of IVIEW-1503 was to decrease the mucociliary clearance using mucoadhesive *in-situ* gel forming formulations to prolong the residence time of the drug in the nasal cavity. Because human nasal mucosa is covered with a layer of approximately 0.1 ml of mucus, which consists of sodium, potassium and calcium ions, a solution-gel phase transition will occur with ion-sensitive gel formulations.

PVP-I is a low-cost topical medication which could significantly reduce the burden on the existing health care system if it is proven to be effective in reducing viral load. Presently, the estimated cost of managing a patient with proven positivity for COVID 19 but without any complications is $9,673. For a patient with complications or co-morbidity it is $13,767 and for a patient with major complications or co-morbidity it increases to $20,292 (54). If we only assess the deceased patients (281,399 by May 10, 2020) and presume that they were hospitalized without complications prior to their death, the amount globally spent to treat this condition would be over 2.72 billion USD. This still does not include testing costs or healthcare costs for those patients with mild to moderate complications.

The next phase is to determine if topical PVP-I nasal sprays and irrigations will result in a reduction in the SARS-CoV-2 viral load, and to show whether there is a significant reduction in hospitalization or need for advanced life support in these patients. Based on a clinical study in Europe, it was found that paucisymptomatic patients, because of high viral loads in upper respiratory tract samples, might potentially transmit the disease during the very first days of infection despite having a mild presentation or symptoms of the disease (55). It therefore seems critical to reduce viral loads at the beginning of the infection, not only for the inhibition of virus evolution for the infected patient, but also for the prevention of virus transmission towards contacts of infected patients.

In addition, it would be important to determine whether there is a significant symptomatic improvement in the quality of life scores with the use of povidone-iodine nasal spray or irrigation. Further development of a long-acting PVP-I gel forming nasal spray, or a PVP-I nasal irrigation formulation, to treat patients infected with COVID-19 in the early stages to lessen the severity of the infection and possibly prevent it from progressing into a severe stage would be important.

## Conclusion

In this study, PVP-I formulations were shown to inactivate SARS-CoV-2 virus efficiently in both dose- and time-dependent manner. This suggests that PVP-I could potentially be used as a disinfectant for SARS-CoV-2 virus. More importantly, IVIEW-1503 PVP-I nasal formulation may potentially be used to inactivate SARS-CoV-2 virus in the nasal cavity thereby preventing infection of the airways. Subsequent development of IVIEW-1201 PVP-I ophthalmic formulation can also possibly be used to inactivate SARS-CoV-2 virus in the eyes and further prevent viral infection. Further human clinical studies to evaluate these sustained release PVP-I gel forming formulations against COVID-19 to assess both safety and efficacy are warranted.

## Acknowledgments

We thank the Institute for Antiviral Research at Utah State University for performing the *in-vitro* virucidal study, IUVO Biosciences in New York and Product Safety Labs in New Jersey for performing *in-vivo* animal toxicology studies for ocular and intranasal administrations, respectively. We thank the University of British Columbia, St. Paul’s Sinus Centre colleagues for help with discussing COVID-19 trials. We also thank NIH NEI SBIR Phase I Grant 1R43EY027238-01 and NIH NIAID Phase I Grant 5R43AI138660 for support of adenoviral conjunctivitis and chronic rhinosinusitis projects, respectively. Funding is provided by IVIEW Therapeutics Inc.

## Disclaimers

Dr. Bo Liang and Dr. John J. Baldwin are co-founders of IVIEW Therapeutics Inc. and have financial interest in the company.

## References

1. “Coronavirus Disease 2019”. World Health Organization. [Cited 2020 May 12]. https://www.who.int/emergencies/diseases/novel-coronavirus-2019

2. “WHO Director-General’s opening remarks at the media briefing on COVID-19-11 March 2020”. World Health Organization. 11 March 2020. [Cited 2020 May 12]. https://www.who.int/dg/speeches/detail/who-director-general-s-opening-remarks-at-the-media-briefing-on-covid-19---11-march-2020

3. Chen J. Pathogenicity and transmissibility of 2019-nCoV-A quick overview and comparison with other emerging viruses. Microbes and Infection. 2020; 22:2:69–71.

4. “Statement on the second meeting of the International Health Regulations (2005) Emergency Committee regarding the outbreak of novel coronavirus (2019-nCoV)”. World Health Organization. 30 January 2020. [Cited 2020 May 12] https://www.who.int/news-room/detail/30-01-2020-statement-on-the-second-meeting-of-the-international-health-regulations-(2005)-emergency-committee-regarding-the-outbreak-of-novel-coronavirus-(2019-ncov).

5. Wang C, Horby PW, Hayden FG, Gao GF. A novel coronavirus outbreak of global health concern. The Lancet. 2020; 395:10223:470–473.

6. Perlman S. Another Decade, Another Coronavirus. New England Journal of Medicine. 2020;382:8:760–62.

7. Cui J, Li F, Shi ZL. Origin and evolution of pathogenic coronaviruses. Nat Rev Microbiol. 2019; 17:3:181–92.

8. De Wit E, van Doremalen N, Falzarano D, Munster VJ. SARS and MERS: Recent insights into emerging coronaviruses. Nat Rev Microbiol. 2016; 14:8:523–34.

9. Lin L, Lu L, Cao W, Li T. Hypothesis for potential pathogenesis of SARS-CoV-2 infection–a review of immune changes in patients with viral pneumonia. Emerging Microbes & Infections. 2020; 9: 1:727–32.

10. Zhou P, Yang XL, Wang XG, Hu B, Zhang L, Si HR, et al. A pneumonia outbreak associated with a new coronavirus of probable bat origin. Nature. 2020; 579:7798:270–73.

11. “Q&A on coronaviruses”. World Health Organization. 11 February 2020. [Cited 2020 May 12]. https://www.who.int/emergencies/diseases/novel-coronavirus-2019/question-and-answers-hub/q-a-detail/q-a-coronaviruses

12. “Q & A on COVID-19”. European Centre for Disease Prevention and Control. 24 April 2020 [Cited 2020 May 12]. https://www.ecdc.europa.eu/en/covid-19/questions-answers

13. “Coronavirus Disease 2019 (COVID-19)—Transmission”. Centers for Disease Control and Prevention. 17 March 2020. [Cited 2020 May 12]. https://www.cdc.gov/coronavirus/2019-ncov/prevent-getting-sick/how-covidspreads.html?CDC_AA_refVal=https%3A%2F%2Fwww.cdc.gov%2Fcoronavirus%2F2019-ncov%2Fprepare%2Ftransmission.html

14. Sungnak W, Huang N, Bécavin C, Berg M, Queen R, Litvinukova M, Talavera-López C, et al, SARS-CoV-2 entry factors are highly expressed in nasal epithelial cells together with innate immune genes. Nature Medicine. 2020; 26: 681–87.

15. Zhou L, Xua Z, Castiglionea GM, Soibermana US, Eberhart CG, Duh EJ. ACE2 and TMPRSS2 are expressed on the human ocular surface, suggesting susceptibility to SARS-CoV-2 infection. BioRxiv. May 9, 2020. https://www.biorxiv.org/content/10.1101/2020.05.09.086165v1

16. DeBroff BM, COVID-19: Ocular manifestations, ocular secretions, and ocular portal of entry. Adv Ophthalmol Vis Syst. 2020; 10:2:48–49.

17. Bhagwat D, Oshlack B. Stabilized PVP-I Solutions. United States Patent #5, 126, 127; 1992 June 30.

18. Liang B, Tessema B, Capriotti JA, Michael SC, Wei S. Pharmaceutical Compositions comprising iodine and steroid and uses thereof for sinus diseases. WO 2012/177251 US 2014/0219949. 2012 Dec 27.

19. Jaya C, Job A, Mathai E, Antonisamy B. Evaluation of topical povidone-iodine in chronic suppurative otitis media. Arch Otolaryngol Head Neck Surg. 2003;129:10:1098–240.

20. Rooijackers-Lemmens E, Van Wijngaarden JJ, Opstelten W, Broen A, Romeijnders ACM. NHG-standard otitis externa. Huisarts Wet. 1995;28:6:265–71.

21. Rowlands S, Devalia H, Smith C, Hubbard R, Dean A. Otitis externa in UK general practice: a survey using the UK General Practice Research Database. Br J Gen Pract. 2001;51:468:533–38.

22. Kavanagh K. Chronic Otitis Externa and Otomycosis. World Articles in Ear Nose & Throat. 27 May 2008. [Cited 2020 May 12] http://www.waent.org/archives/2008/vol1/chronic_otitis_externa/otomycosis.htm

23. Perez R, Freeman S, Sohmer H, Sichel JY. Vestibular and cochlear ototoxicity of topical antiseptics assessed by evoked potentials. Laryngoscope. 2000; 110:9:1522–27.

24. Rezapoor M, Nicholson T, Tabatabaee RM, Chen AF, Maltenfort MG, Parvizi, J. Povidone-Iodine–Based Solutions for Decolonization of Nasal Staphylococcus Aureus: A Randomized, Prospective, Placebo-Controlled Study. The Journal of Arthroplasty. 2017; 32:9:2815–19.

25. Anderson MJ, David ML, Scholz M, Bull SJ, Morse D, Hulse-Stevens M, et al. Efficacy of Skin and Nasal Povidone-Iodine Preparation against Mupirocin-Resistant Methicillin-Resistant Staphylococcus Aureus and S. Aureus within the Anterior Nares. Antimicrob Agents Chemother. 2015;59:5:2765–73.

26. Peng HM, Wang LC, Zhai JL, Weng XS, Feng B, Wang W. Effectiveness of Preoperative Decolonization with Nasal Povidone Iodine in Chinese Patients Undergoing Elective Orthopedic Surgery: A Prospective Cross-Sectional Study. Braz J Med Biol Res. 2017;51:2:e6736

27. Phillips M, Rosenberg A, Shopsin B, Cuff G, Skeete F, Foti A, et al. Preventing Surgical Site Infections: A Randomized, Open-Label Trial of Nasal Mupirocin Ointment and Nasal Povidone-Iodine Solution. Infect Control Hosp Epidemiol. 2014; 35:7:826–32.

28. Gluck U, Martin U, Bosse B, Reimer K, Mueller S. A Clinical Study on the Tolerability of a Liposomal Povidone-Iodine Nasal Spray: Implications for Further Development. ORL J Otorhinolaryngol Relat Spec. 2007; 69:2:92–99.

29. Wood A, Payne D. The action of three antiseptics/disinfectants against enveloped and non-enveloped viruses. J Hosp Infect. 1998; 38:4:283–95.

30. Kanagalingam J, Feliciano R, Hah JH, Labib H, Le TA, Lin JC. Practical use of povidone-iodine antiseptic in the maintenance of oral health and in the prevention and treatment of common oropharyngeal infections. Int J Clin Pract. 2015; 69:11:1247–56.

31. Sauerbrei A, Wutzler P. Virucidal efficacy of povidone-iodine-containing disinfectants. Lett Appl Microbiol. 2010; 51:2:158–63.

32. Wada H, Nojima Y, Ogawa S, Hayashi N, Sugiyama N, Kajiura T, et al. Relationship between Virucidal Efficacy and Free Iodine Concentration of Povidone-Iodine in Buffer Solution. Biocontrol Sci. 2016; 21:1:21–7.

33. Kariwa H, Fujii N, Takashima I. Inactivation of SARS coronavirus by means of povidone-iodine, physical conditions and chemical reagents. Dermatology. 2006; 212:1:119–23.

34. Eggers M, Koburger-Janssen T, Eickmann M, Zorn J. In Vitro Bactericidal and Virucidal Efficacy of Povidone-Iodine Gargle/Mouthwash Against Respiratory and Oral Tract Pathogens. Infect Dis Ther. 2018; 7:2:249–59.

35. Eggers M, Eickmann M, Zorn J. Rapid and Effective Virucidal Activity of Povidone-Iodine Products Against Middle East Respiratory Syndrome Coronavirus (MERS-CoV) and Modified Vaccinia Virus Ankara (MVA). Infect Dis Ther. 2015; 4:4:491–501.

36. Eggers M, Koburger-Janssen T, Ward LS, Newby C, Müller S. Bactericidal and Virucidal Activity of Povidone-Iodine and Chlorhexidine Gluconate Cleansers in an In Vivo Hand Hygiene Clinical Simulation Study. Infect Dis Ther. 2018; 7:2:235–247.

37. Matsuhira T, Kaji C, Murakami S, Maebashi K, Oka T, Takeda N, et al. Evaluation of four antiseptics using a novel murine norovirus. Exp Anim. 2012; 61:1:35–40.

38. Eggers M, Eickmann M, Kowalski K, Zorn J, Reimer K. Povidone-iodine hand wash and hand rub products demonstrated excellent in vitro virucidal efficacy against Ebola virus and modified vaccinia virus Ankara, the new European test virus for enveloped viruses. BMC Infect Dis. 2015; 15:375.

39. Kirk-Bayley J, Combes J, Sunkaraneni S, Challacombe S. The Use of Povidone Iodine Nasal Spray and Mouthwash During the Current COVID-19 Pandemic May Reduce Cross Infection and Protect Healthcare Workers. SSRN. https://ssrn.com/abstract=3563092

40. Lee VHL, Robinson JR. Mechanistic and quantitative evaluation of precorneal pilocarpine disposition in albino rabbits. J Pharm Sci. 1979; 68:6:673–84.

41. Liang, B., Baldwin J., Wei G. Pharmaceutical Formulations that Form Gel In Situ. WO2017/074965 and US2017/0266294. 2017 April 5.

42. Liang, B. In Situ Gel-Forming Pharmaceutical Compositions and Uses Thereof For Sinus Diseases. WO2019/046844. 2019 July 3.

43. Balasubramaniam J, Kant S, Pandit JK. In vitro and in vivo evaluation of the Gelrite gellan gum-based ocular delivery system for indomethacin. Acta Pharm. 2003; 53:4:251–61.

44. Chu CM, Cheng VC, Hung IF, Wong MM, Chan KH, Chan KS, et al. Role of lopinavir/ritonavir in the treatment of SARS: Initial virological and clinical findings. Thorax. 2004; 59:3:252–256.

45. Savarino A, Di Trani L, Donatelli I, Cauda R, Cassone A. New insights into the antiviral effects of chloroquine. Lancet Infect Dis. 2006; 6:2:67–9.

46. Wang M, Cao R, Zhang L, Yang X, Liu J, Xu M, et al. Remdesivir and chloroquine effectively inhibit the recently emerged novel coronavirus (2019-nCoV) in vitro. Cell Res. 2020; 30:3:269–71

47. “Researchers find two new drugs that can effectively inhibit coronavirus” 4 February 2020. [Cited 2020 May 12]. https://news.cgtn.com/news/2020-02-04/Researchers-find-two-drugs-that-can-effectively-inhibit-coronavirus-NOFpci7NJK/index.html

48. Dong L, Hu S, Gao J. Discovering drugs to treat coronavirus disease 2019 (COVID-19). Drug Discov Ther. 2020; 14:1:58–60.

49. Stahlmann R, Lode H. Medication for COVID-19-an Overview of Approaches Currently Under Study. Dtsch Arztebl Int. 2020; 117:13:213–19.

50. Magro G. SARS-CoV-2 and COVID-19: What are our options? Where should we focus our attention on to find new drugs and strategies? Travel Med Infect Dis. 2020:101685.

51. Mehta N, Mazer-Amirshahi M, Alkindi N, Pourmand A. Pharmacotherapy in COVID-19; A narrative review for emergency providers. Am J Emerg Med. 2020; http:https://www.ajemjournal.com/article/S0735-6757(20)30263-1/pdf

52. Augustin M, Hallek M, Nitschmann S. Remdesivir in patients with severe COVID-19. Internist (Berl). 2020. https://link.springer.com/article/10.1007/s00108-020-00800-5

53. Grein J, Ohmagari N, Shin D, Diaz G, Asperges E, Castagna A, et al. Compassionate use of remdesivir for patients with severe Covid-19. N Engl J Med. 2020; https://www.nejm.org/doi/10.1056/NEJMoa2007016

54. Rae M, Claxon G, Kurani N, McDermott D, Cox C. Potential Costs of Coronavirus Treatment for People with Employer Coverage. Peterson-Kaiser Health System Tracker, 13 Mar. 2020. [Cited 2020 May 12]. www.healthsystemtracker.org/brief/potential-costs-of-coronavirus-treatment-for-people-with-employer-coverage/

55. Lescure FX, Bouadma L, Nguyen D, Parisey M, Wickcy PH, Behillil S, et al. Clinical and virological data of the first cases of COVID-19 in Europe: a case series. The Lancet Infectious Diseases. 2020. https://www.thelancet.com/journals/laninf/article/PIIS1473-3099(20)30200-0/fulltext

